## Supplementary figures and images for "The second messenger signaling molecule cyclic di-AMP drives developmental cycle progression in *Chlamydia trachomatis*"

### Supplemental Figure 1

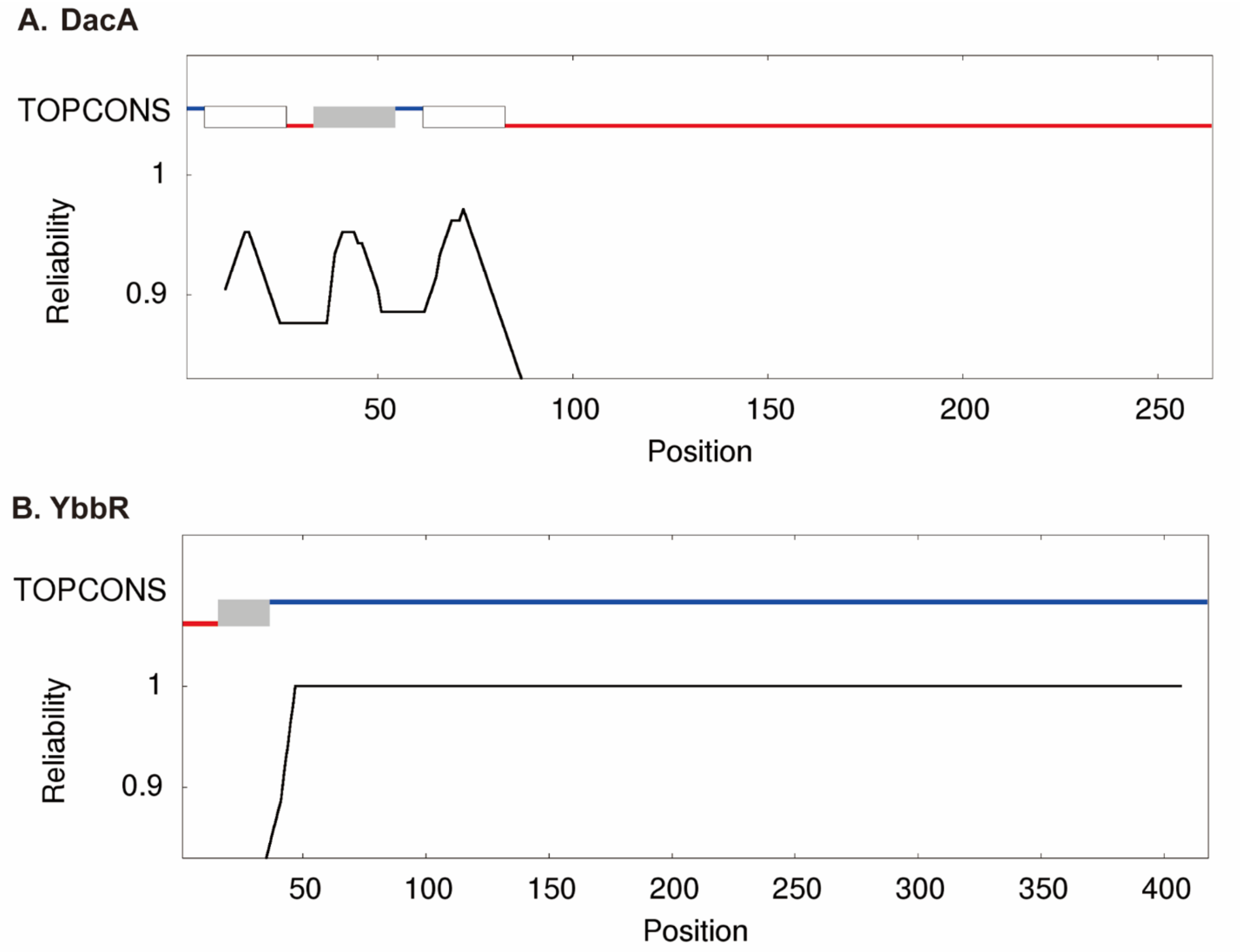

### Supplemental Figure 2

Fig.S2

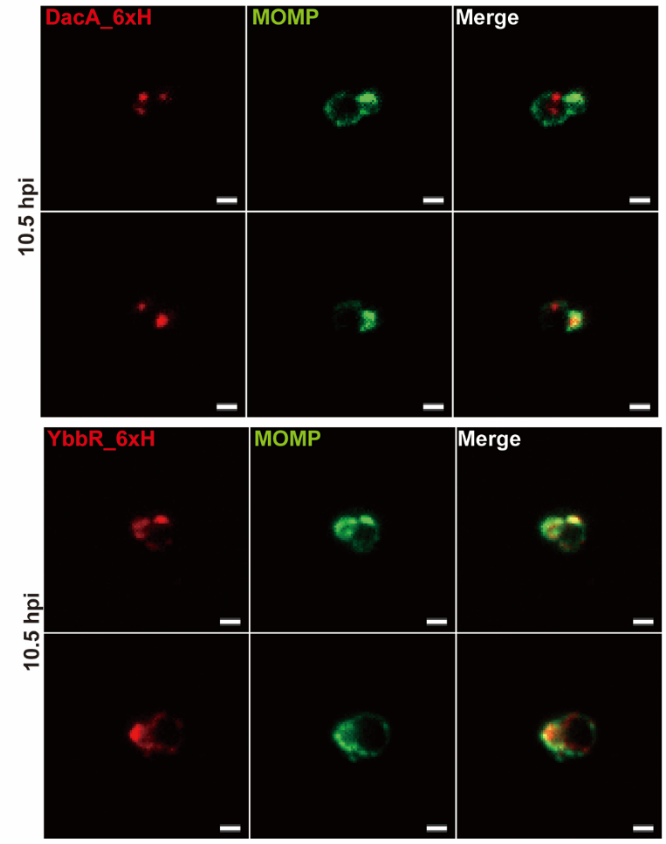

### Supplemental Figure 3

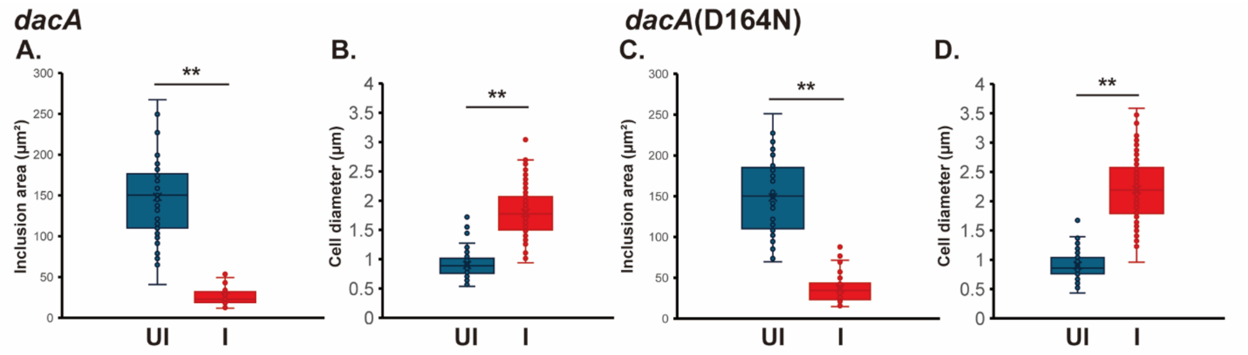

### Supplemental Figure 4

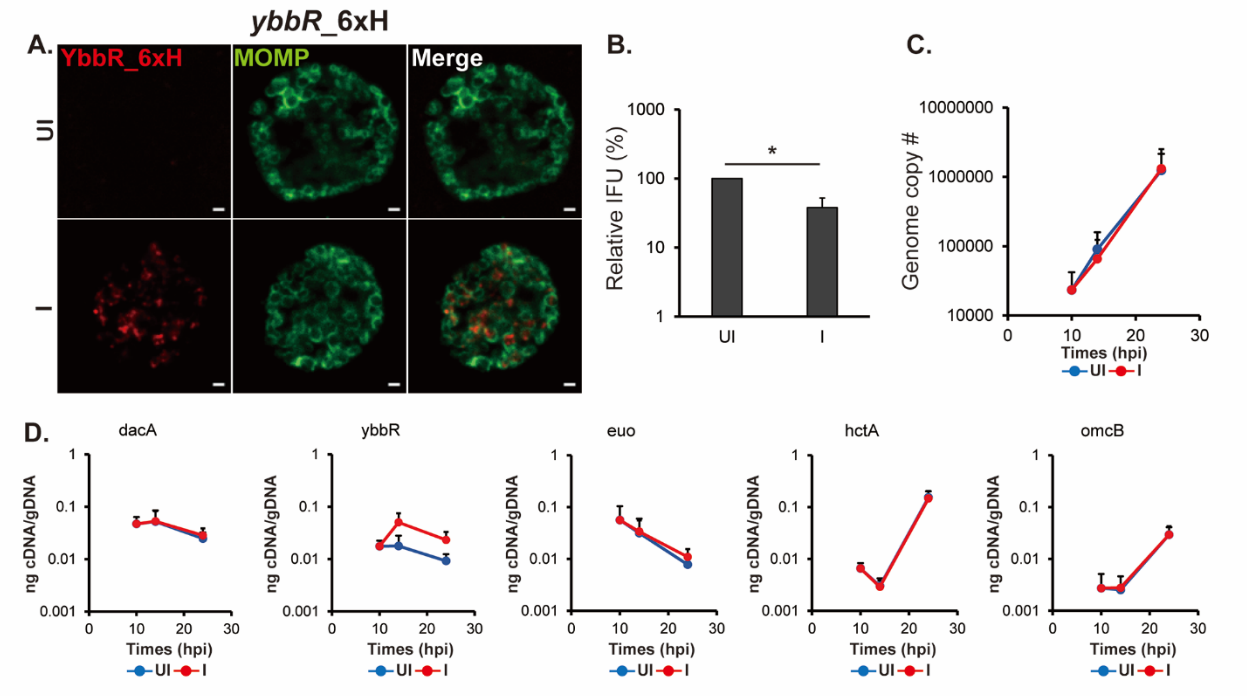

### Supplemental Figure 5

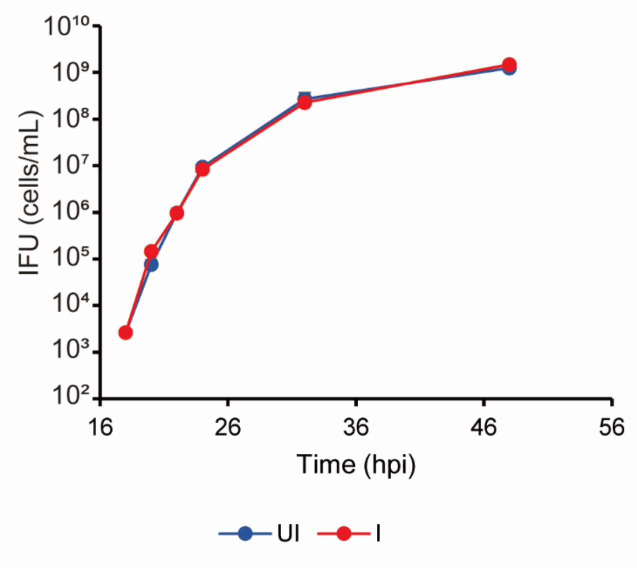

### Supplemental Figure 6

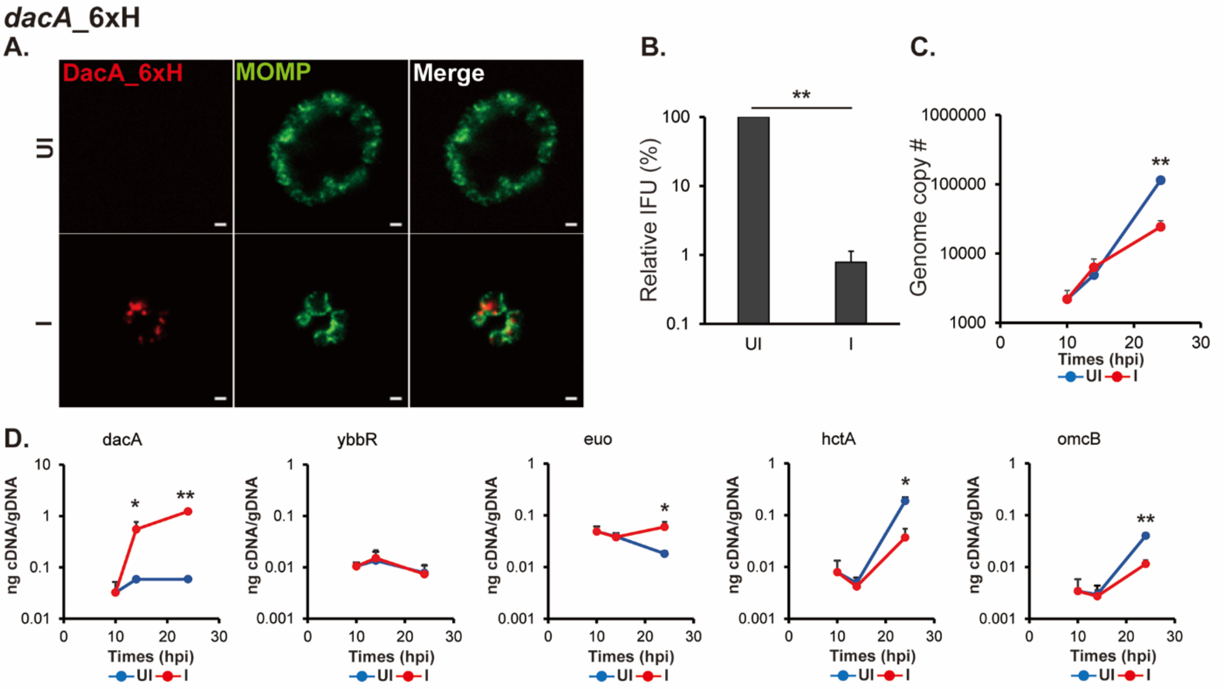

### Supplemental Figure 7

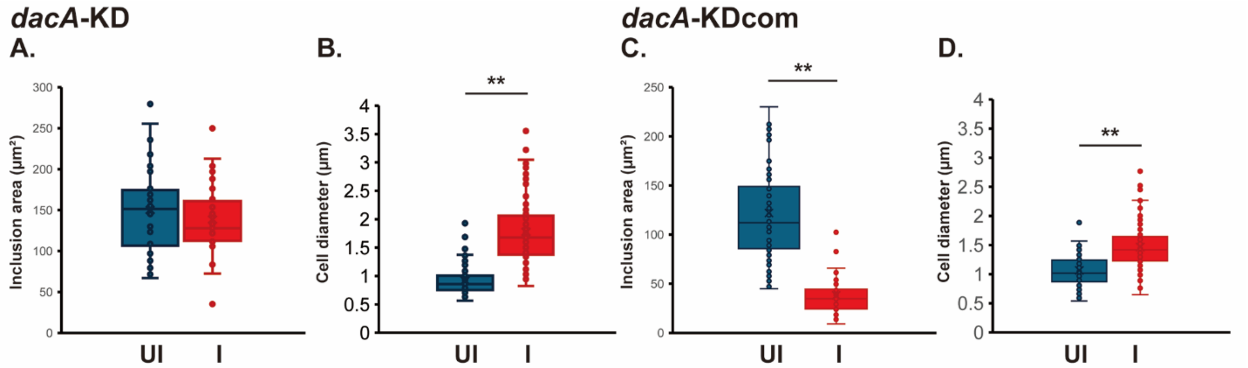

### Supplemental Figure 8

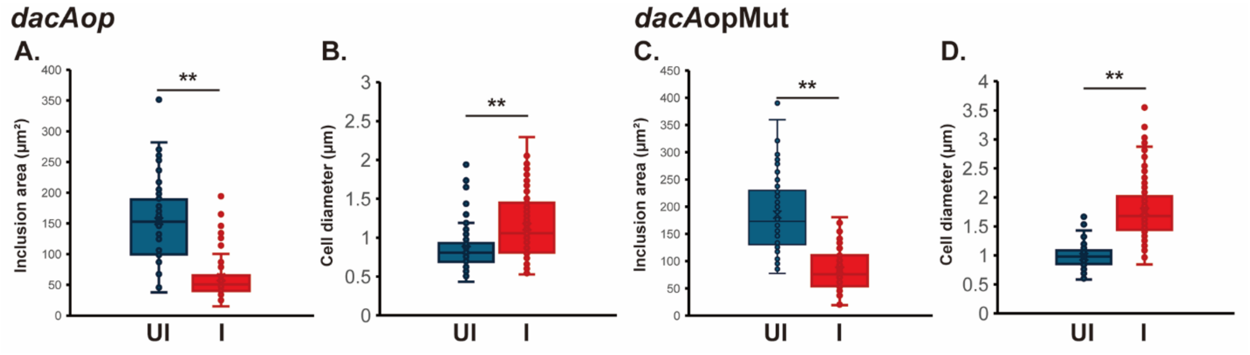

### Supplemental Figure 9

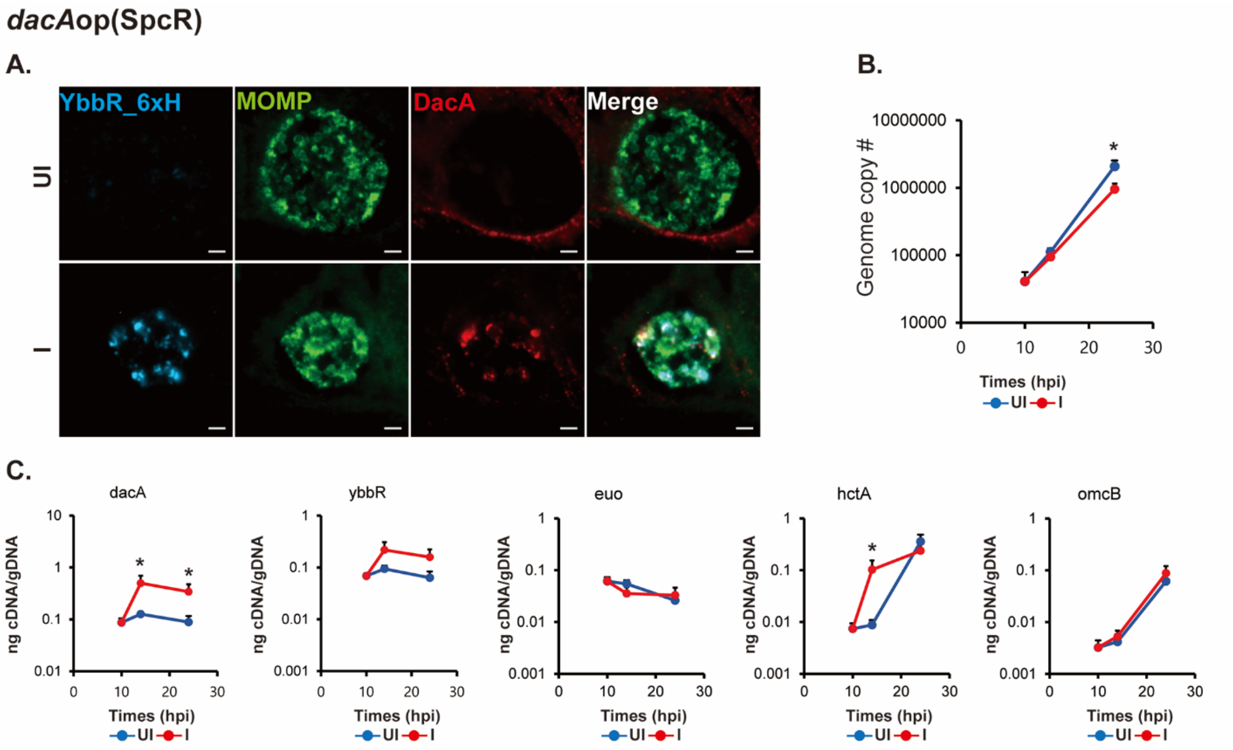

### Supplemental Figure 10

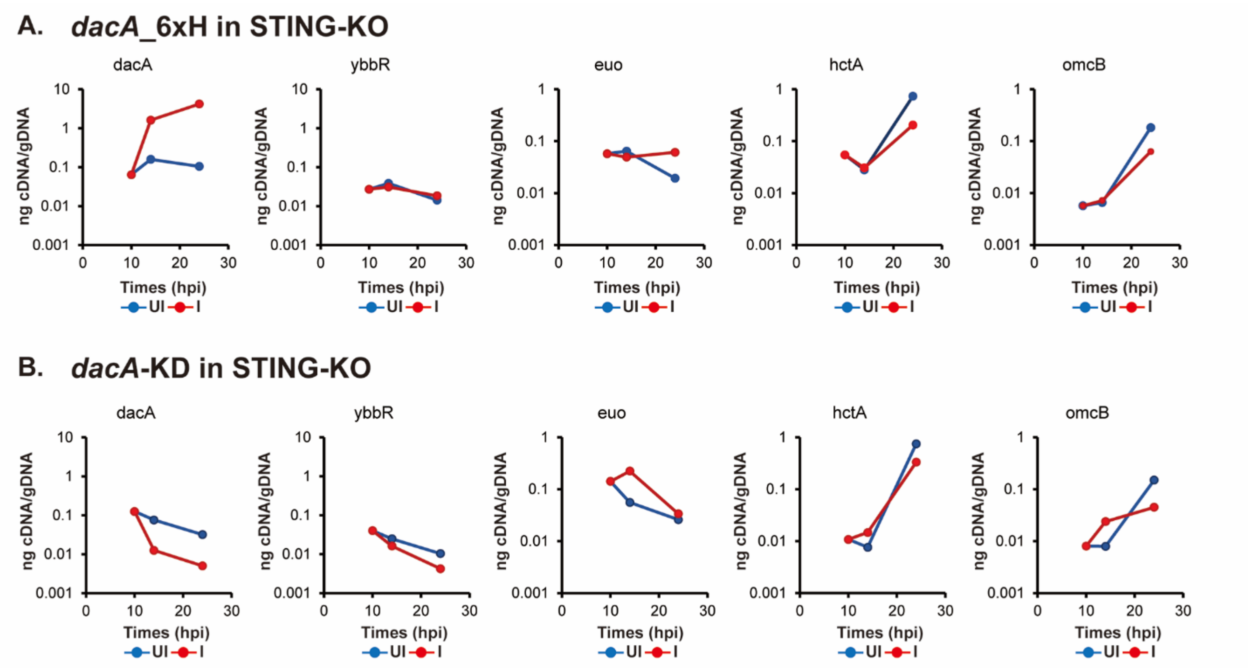
