## Supplemental Table S1 for "The second messenger signaling molecule cyclic di-AMP drives developmental cycle progression in *Chlamydia trachomatis*"

**Supplementary Table 1. List of Plasmids and Primers used in the study**

| **Construct Plasmid** | **Relevant genotype** | **Ori** | **Source of Reference** |
| --- | --- | --- | --- |
| pBOMBL | *bla* P*tet*::*mCherry* P*_Nm_*::*gfp* | pUC19 | (1, 2) |
| pBOMBL-spc^r^ | *aadA P_tet_::mCherry* P*_Nm_*::*gfp* | pUC19 | This study |
| pBOMBL-*dacA*_6xH | *bla* P*tet*::Ctr_*dacA _6xH* P*_Nm_*::*gfp* | pUC19 | This study |
| pBOMBL- *dacA* | *bla* P*tet*::Ctr_*dacA* P*_Nm_*::*gfp* | pUC19 | This study |
| pBOMBL- *dacA*(D164N) | *bla* P*tet*::Ctr_*dacA*(D164N) P*_Nm_*::*gfp* | pUC19 | This study |
| pBOMBL- *ybbR*_6xH | *bla* P*tet*::Ctr_ *ybbR*_6xH P*_Nm_*::*gfp* | pUC19 | This study |
| pBOMBL-*dacA-ybbR*_6xH | *bla* P*tet*::Ctr_*dacA-ybbR*_6xH P*_Nm_*::*gfp* | pUC19 | This study |
| pBOMBL-spc^r^-*dacA-ybbR*_6xH | *aadA P_tet_::*Ctr*_dacA-ybbR_*6xH P*_Nm_*::*gfp* | pUC19 | This study |
| pBOMBL-*dacA(D164N)-ybbR*_6xH | *bla* P*tet*::Ctr_*dacA*(D164N)*-ybbR_6xH* P*_Nm_*::*gfp* | pUC19 | This study |
| pL12CRia(*dacA*-IGR) | *bla* P*tet*::*ddAsCpf1*vaa P*_dnaKmut_*::As_crRNA (*dacA*) P*_Nm_*::*gfp* | pUC19 | This study |
| pL12CRia(*dacA*-IGR)-*dacA-ybbR*_6xH | *bla* P*tet*::*ddAsCpf1vaa*-*dacA*-*ybbR*_6xH P*_dnaKmut_*::As_crRNA (*dacA*) P*_Nm_*::*gfp* | pUC19 | This study |
| pBOMBLv2-ΔTM*dacA* | *aadA P_tet_::*Ctr*_*ΔTM*dacA* P*_Nm_*::*gfp* | pUC19 | This study |
| pBOMBLv2-ΔTM*dacA*(D164N) | *aadA P_tet_::*Ctr*_*ΔTM*dacA*(D164N) P*_Nm_*::*gfp* | pUC19 | This study |

| **Primer name** | **Sequence** | **Features** | **Usage** |
| --- | --- | --- | --- |
| dacA/(pBOMBL)/F | aaagatcttcacacaggacatctgcATGTTCGTAGGTATAACGTATTACAC | lower case for plasmid overlap construction | for amplification of *dacA+/-6xH and dacA-ybbR_6xH* |
| dacA_6xH/(pBOMB)/R | acatatttgaatggtcgaccggtacttaatggtgatggtgatggtgTTTTTTACGCATCCAGGAG | lower case for plasmid overlap construction and lower case with underline for 6xHis tag | for amplification of *dacA_6xH* |
| dacA/pBOMBL/R | tttgaatggtcgaccggtacTCATTTTTTACGCATCCAGG | lower case for plasmid overlap construction | for amplification of *dacA* without a tag |
| ybbR/(pBOMBL)/F | aaagatcttcacacaggacatctgcATGATCAATTTCGTCTCTTG | lower case for plasmid overlap construction | for amplification of *ybbR_6xH* |
| ybbR_6xH/(pBOMB)/R | acatatttgaatggtcgaccggtacttaatggtgatggtgatggtgGGAAGATTTTTTCTTTAGAGAGG | lower case for plasmid overlap construction and 6xHis tag | for amplification of *ybbR_6xH and dacA-ybbR_6xH* |
| dacA/(dCas12)/F | cgtagctgcttaagtaccgg*aggagaatctgc*ATGTTCGTAGGTATAACGTATTAC | lower case for plasmid overlap construction, lower case and underlined with italics for RBS site | For amplification of *dacA-ybbR_6xH* to complement *dacA*-KD |
| dacA87/(pBOMBL)/F | tcttcacacaggacatctgcATGTCTAGGATACGCTTGC | lower case for plasmid overlap construction | for amplification of ΔTM*dacA* |
| ct012_dacA qPCR_F | CGCTTGCGTAGAGGGAAAT | qPCR primer | Amplification of *dacA* |
| ct012_dacA qPCR_R | AGCTCCGATTTGTCGTTCAG | qPCR primer | Amplification of *dacA* |
| ct011_ybbR qPCR_F | GCTTCGCCCTTCCAATCT | qPCR primer | Amplification of *ybbR* |
| ct011_ybbR qPCR_R | GATGTCTGTGTCCAAGCATACTA | qPCR primer | Amplification of *ybbR* |
| hctA qPCR F | AAGCTAAAGCTGCTGCTAAGA | qPCR primer | Amplification of *hctA* |
| hctA qPCR R | GTTGGTTTGACCTTTGCTTTAGT | qPCR primer | Amplification of *hctA* |
| omcB qPCR F | CGGTAGGATCTCCCTATCCTATT | qPCR primer | Amplification of *omcB_359-457nt_* |
| omcB qPCR R | CGAACTCTGCTTCACATGGTA | qPCR primer | Amplification of *omcB_359-457nt_* |
| euo qPCR F | CGAAGACTACTCGTTGGGAAATA | qPCR primer | Amplification of *euo_179-278nt_* |
| euo qPCR R | AACAGAAGCTCTCCTTGATAAGT | qPCR primer | Amplification of *euo_179-278nt_* |

| **gBlock name** | **Sequence** | **Features** | **Usage** |
| --- | --- | --- | --- |
| *dacA* crRNA | tgtgaaagtgggtcttaagacgtcggtactgcatgtgacgcacgtagatcatgca*TTCACCGGTGGAGACGGTTTTCTTATAATGACACC*TAATTTCTACTCTTGTAGAT**CTGAAAATAATGAAAACACTT**CAAATAAAACGAAAGGCTCAGTCGAAAGACTGGGCCTTTCGTTTTATcaacagcggtctactgaatctgagctagtgcgtgatataattaaaattatattca | Lower case for plasmid overlap and spacer, *italicized* for P_dnaKmut_ promoter sequence, underlined for crRNA scaffold, **bold** for *dacA* targeting sequence, Upper case for rrnB1 terminator | For CRISPRi knockdown of *dacA* operon |

1.	Bauler LD, Hackstadt T. 2014. Expression and targeting of secreted proteins from Chlamydia trachomatis. J Bacteriol 196:1325-34.

2.	Ouellette SP, Blay EA, Hatch ND, Fisher-Marvin LA. 2021. CRISPR Interference To Inducibly Repress Gene Expression in Chlamydia trachomatis. Infect Immun 89:e0010821.
