## Supplemental Figure Legends for "The second messenger signaling molecule cyclic di-AMP drives developmental cycle progression in *Chlamydia trachomatis*"

**Lee and Ouellette Supplemental Figure Legends**

**Figure S1. The predicted transmembrane domains in DacA and YbbR.** Transmembrane domains in DacA (A) and YbbR (B) were predicted with TOPCONS (https://topcons.cbr.su.se/)(55). The red and blue lines represent cytosolic and periplasmic domains, respectively.

**Figure S2.** **The localization of DacA_6xH (A) and YbbR_6xH (B) in the first dividing cells.** *C. trachomatis* encoding DacA_6xH or YbbR_6xH was infected into HeLa cells. At 4 hpi, the constructs were induced with 5 nM aTc. At 10.5 hpi, the infected cells were fixed with an aldehyde fixing solution (3.2% Formaldehyde, 0.022% Glutaraldehyde in 1X PBS) for 2 min and permeabilized with 90% MeOH for 1 min. The images were acquired on a Zeiss Imager.Z2 equipped with an Apotome2 using a 100X lens objective.

**Figure S3. Inclusion area and cell diameter measurements of *dacA* and *dacA*(D164N)*.*** **A.** Inclusion area of the *dacA* strain. We measured the area of 56 inclusions (from n=3 replicates). **B.** Cell diameter of the *dacA* strain. We measured the diameter of 142 (uninduced = UI) or 119 (induced = I) bacteria (from n=3 replicates). **C.** Inclusion area of the *dacA*(D164N) strain. We measured the area of 54 inclusions (from n=3 replicates). **D.** Cell diameter of the *dacA*(D164N) strain. We measured the diameter of 143 (UI) or 145 (I) bacteria (from n=3 replicates). The inclusion area and cell diameter were measured using Fiji software. **: p<0.001 via two sample equal variance t-test.

**Figure S4. Overexpression of YbbR_6xH does not affect the chlamydial developmental cycle.** HeLa cells were infected with *C. trachomatis* transformed with a plasmid encoding an anhydrotetracycline-inducible YbbR_6xH (i.e., *ybbR*_6xH; see Fig. 1D). At 10 hpi, expression of the construct was induced or not with 5 nM aTc, and IFA, IFU, DNA, and RNA samples were collected. **A.** IFA images of *ybbR*_6xH strain at 24 hpi. **B.** Quantification of IFU from uninduced and induced samples at 24 hpi. **C.** Quantification of genomic DNA copy number by qPCR in uninduced and induced samples. **D.** Quantification of transcripts by RT-qPCR for *dacA,* *ybbR*, *euo*, *hctA*, and *omcB.* IFA images were acquired on a Zeiss AxioImager.Z2 equipped with an Apotome2 using a 100X lens objective. Scale bar: 1 µm. UI = uninduced (i.e., -aTc); I = induced (i.e., +aTc) for all sample types. *: p<0.05 via two sample equal variance t-test.

**Figure S5. Overexpressed mCherry from the vector control does not affect EB progeny production during the chlamydial developmental cycle.** HeLa cells were infected with *C. trachomatis* transformed with a plasmid encoding an anhydrotetracycline-inducible mCherry. At 10 hpi, expression of the construct was induced or not with 5 nM aTc, and the EB samples were collected at 18, 20, 22, 24, 32, and 48 hpi.

**Figure S6. Overexpression of DacA_6xH is detrimental to the chlamydial developmental cycle.** HeLa cells were infected with *C. trachomatis* transformed with a plasmid encoding an anhydrotetracycline (aTc)-inducible DacA_6xH. At 10 hpi, expression of the construct was induced or not with 5 nM aTc, and DNA and RNA samples were collected at 10, 14, and 24 hpi. Immunofluorescence analysis (IFA) and inclusion forming units (IFU) samples were collected at 24hpi. **A.** IFA images of the *dacA*_6xH strain at 24 hpi. Shown are individual panels for DacA_6xH and major outer membrane protein (MOMP) labeling as well as the merged image. IFA images were acquired on a Zeiss AxioImager.Z2 equipped with an Apotome2 using a 100X lens objective. Scale bar: 1 µm. **B.** Quantification of IFUs from uninduced and induced samples at 24 hpi. **C.** Quantification of genomic DNA copy number by qPCR in uninduced and induced samples. **D.** Quantification of transcripts by RT-qPCR for *dacA,* *ybbR*, *euo*, *hctA*, and *omcB* from uninduced and induced samples*.* UI = uninduced (i.e., -aTc); I = induced (i.e., +aTc) for all sample types. *: p<0.05; **: p<0.001 via two sample equal variance t-test.

**Figure S7. Inclusion area and cell diameter measurements of *dacA*-KD and *dacA*-KDcom.** **A.** Inclusion area of the *dacA*-KD strain. We measured the area of 30 (uninduced = UI) and 29 (induced = I) inclusions (from n=3 replicates). **B.** Cell diameter of the *dacA*-KD strain. We measured the diameter of 126 (UI) or 129 (induced = I) bacteria (from n=3 replicates). **C.** Inclusion area of the *dacA*-KDcom strain. We measured the area of 62 (UI) and 60 (I) inclusions (from n=3 replicates). **D.** Cell diameter of the *dacA*-KDcom strain. We measured the diameter of 149 (UI) and 142 (I) bacteria (from n=3 replicates). The inclusion area and cell diameter were measured using Fiji software. **: p<0.001 via two sample equal variance t-test.

**Figure S8. Inclusion area and cell diameter measurements of *dacA*op and *dacA*opMut.** **A.** Inclusion area of the *dacA*op strain. We measured the area of 52 (uninduced = UI) and 58 (induced = I) inclusions (from n=3 replicates). **B.** Cell diameter of the *dacA*op strain. We measured the diameter of 125 (UI) or 131 (induced = I) bacteria (from n=3 replicates). **C.** Inclusion area of the *dacA*opMut strain. We measured the area of 50 (UI) and 52 (I) inclusions (from n=3 replicates). **D.** Cell diameter of the *dacA*opMut strain. We measured the diameter of 124 (UI) and 150 (I) bacteria (from n=3 replicates). The inclusion area and cell diameter were measured using Fiji software. **: p<0.001 via two sample equal variance t-test.

**Figure S9. Phenotypic characterization of a *dacA*op overexpression construct encoding spectinomycin resistance (Spc^R^).** HeLa cells were infected with *C. trachomatis* transformed with a *dacA*op overexpression construct encoding Spc^R^. At 10 hpi, expression of the construct was induced or not with 5 nM aTc, and DNA and RNA samples were collected at 10, 14, and 24 hpi. Immunofluorescence analysis (IFA) samples were collected at 24hpi. **A.** IFA images of the *dacA*op strain at 24 hpi. Shown are individual panels for DacA, YbbR_6xH, and major outer membrane protein (MOMP) labeling as well as the merged image. **B.** Quantification of genomic DNA copy number by qPCR in uninduced and induced samples. **C.** Quantification of transcripts by RT-qPCR for *dacA, ybbR, euo, hctA,* and *omcB*. IFA images were acquired on a Zeiss AxioImager.Z2 equipped with an Apotome2 using a 100X lens objective. Scale bar: 2 µm. UI = uninduced (i.e., -aTc); I = induced (i.e., +aTc) for all sample types. *: p<0.05 via two sample equal variance t-test.

**Figure S10.** **Quantification of transcripts by RT-qPCR for *dacA,* *ybbR*, *euo*, *hctA*, and *omcB* from *dacA*-KD and *dacA*_6xH strains cultured in STING-KO HeLa cells*.*** STING-KO HeLa cells were infected with *C. trachomatis* transformed with a plasmid encoding an anhydrotetracycline (aTc)-inducible **A.** DacA_6xH and/or **B.** CRISPRi-dCas12 system targeting the *dacA* promoter. At 10 hpi, expression of the construct was induced or not with 5 nM aTc, and DNA and RNA samples were collected at 10, 14, and 24 hpi.
